# Highly oxidized albumin is mainly cleared by mouse liver sinusoidal endothelial cells via the receptors stabilin-1 and -2

**DOI:** 10.1101/2023.07.20.549630

**Authors:** Christopher F. Holte, Karolina J. Szafranska, Larissa D. Kruse, Jaione Simon-Santamaria, Ruomei Li, Dmitri Svistounov, Peter A.G. McCourt

## Abstract

**Background:** Oxidized albumin (oxHSA) is elevated in several pathological conditions, especially those involving the liver, such as decompensated cirrhosis, acute on chronic liver failure and liver mediated renal failure. Patient derived oxidized albumin was previously shown to be an inflammatory mediator in cultured endothelial cells and leukocytes. The removal from circulation of oxidized albumins is therefore essential for maintenance of homeostasis. Normal serum levels of oxidized albumin are low, implying it is constantly eliminated. Liver sinusoidal endothelial cells (LSEC) are prominent scavenger cells in the body, specializing in the removal of macromolecules e.g. hyaluronan, denatured collagen, modified albumins, bacterial endotoxin (LPS) and oxidized lipoprotein. Given that oxidized albumin is mainly cleared by the liver, we hypothesize the LSEC are the site of uptake in the liver. Furthermore the stabilins -1 and -2 are the most prominent candidates for oxHSA uptake receptors, given their expression pattern and uptake of other ligands.

**Methods:** In vivo biodistribution, hepatocellular distribution and in vitro uptake studies on isolated liver cell populations or receptor expressing cell lines.

**Results:** In vivo oxHSA was cleared rapidly (t1/2 <90seconds) by the liver (47% of uptake) and distributed to mainly the LSEC. In in vitro studies LSEC endocytosed oxHSA much more than other cell populations isolated from the liver. Furthermore, it was shown that the uptake was mediated by the stabilins, by inhibiting uptake in LSEC with other stabilin ligands and showing uptake in HEK cells overexpressing stabilin-1 or 2. oxHSA also inhibited the uptake of other stabilin ligands.

**Conclusions:** LSEC and their stabilins are vital for the clearance of oxidized albumin, and therefore play a pivotal role in maintaining homeostasis.

## Introduction

Albumin is the most abundant protein in blood (40g/L of plasma is made up of albumin^1^, and it has a correspondingly large number of functions, including binding and transporting a host of ligands including but not limited to free fatty acids, drugs (including: warfarin, salicyclic acid, propofol, lidocaine) and metabolites ^2^. In the blood stream albumin serves as the main antioxidant due to its readily reacting cysteine-34 residue and metal ion binding properties, and is also found extensively in the extravascular extracellular space ^3,2^. Albumin is therefore a vital component in mitigation of oxidative stress throughout the body.

Oxidative stress is implicated in the pathophysiology of several diseases, such as atherosclerosis; where oxidation products are linked with plaque formation ^4^; nephrotic damage in leukocyte-dependent glomerulonephritis ^5^; and the development and progression of neurodegenerative diseases ^6^. Ischemia-modified albumin, thought to be formed by reaction with reactive oxygen species and or hydroxyl radicals ^7^, therefore a form of oxidized albumin, is a marker of poor prognosis in patients reporting chest pain, and is determined clinically by assaying the cobalt binding ability of patient sera ^8^. The neutrophil myeloperoxidase is one endogenous system capable of producing extremely potent oxidants such as hypochlorite, thio- and hypothiocyanite ^9^, thus serving as a link between inflammation and oxidative stress.

Oxidative stress and the presence of oxidized albumin is also a component of the pathogenesis of acute on chronic liver failure ^10,11^, a syndrome that develops from decompensated cirrhosis ^12^. Elevated advanced oxidation protein products (AOPP) and modified albumins were also found in plasma samples from idiosyncratic drug-induced liver injury ^13^. Oxidation of serum albumin in patients with cirrhosis and bacterial peritonitis causes decreased binding properties of albumin, predicting impaired transport function ^14^. *In vitro* oxidized albumin has been shown to have altered affinities, both increased and decreased, to various drugs and metabolites ^15, 16^.

Oxidised albumin is associated with a number of other pathologies. There is a correlation between the fraction of oxidized albumin and atherosclerosis development ^17^. In nephrotic patients oxidized and advanced glycation end-product (AGE) albumin was found, and a reduction in oxidized albumin considered a beneficial marker after hemodialysis ^18^. The oxidation products themselves have been suggested to be uremic toxins playing an active role in the development of chronic renal failure ^19^. An increased fraction of oxidized relative vs. non-oxidized albumin is also characteristic of Diabetes Mellitus patients ^20^. Oxidized albumin from hypoalbuminemic hemodialysis patient samples were shown to cause elevated expression of inflammatory cytokines in HUVECs ^21^ and primary peripheral blood leukocytes ^22^. The proinflammatory effect was shown to be oxidation dependent and reversible upon chemical reduction of the albumin ^21^. Oxidative modifications of HSA have been shown to induce clearance from circulation ^23^, showing a potential link between hypoalbuminemia and oxidative stress often observed in cirrhosis. Iwao ^24^ found chemically oxidized HSA (oxHSA), produced using the hypochlorite analogue chloramine-T, to be similar to oxidized albumin found in uremic patients. This oxHSA was found to be rapidly cleared from circulation in mice, primarily by the liver (51%) and spleen (23%), which are two of the major scavenging organs in the body. Liver sinusoidal endothelial cells (LSEC) are known to take up a host of macromolecular waste from the bloodstream ^25^ whereas Kupffer cells (KC), the liver resident macrophages, remove larger (>200 nm) complexes from the circulation ^26^. Modified albumins such as Advanced Glycation End-products-BSA (AGE-BSA) and formaldehyde modified BSA (FSA) are taken up by the liver sinusoidal endothelium, ^27,28,29^ the scavenging endothelium of the liver sinusoids. AGEs ^30,31^ FSA ^32,33,27^ were shown to be primarily endocytosed via the scavenger receptor class H ^34^ (SR-H), also known as stabilin-1 and -2. Oxidized low density lipoproteins oxLDL ^35^ and acetylated LDL ^36^ were also shown to be taken up by the liver sinusoidal endothelial cells via stabilin-1 and -2. Stabilin-1/2 double knockout mice exhibit glomerular fibrosis, with significant reduction to the animal lifespan, indicating that a reduction in clearance via the stabilins in the liver had downstream effects on the kidneys ^37^. The liver and spleen are the main sites of stabilin 1 and 2 expression ^38,39^.

The ability to induce oxidative stress-like damage of oxidation protein products combined with the rapid uptake of oxHSA by the liver and the propensity for LSEC to clear modified proteins, make these cells a potential site of clearance of oxidized albumin and of injury during sustained oxidative stress.

If oxHSA binds to a scavenger receptor, as its rapid clearance suggests, this may have implications for the clearance of other waste molecules by the same. We therefore sought to determine which cell type and receptor take up oxHSA in the liver and describe the effects of oxHSA upon these cells.

## Experimental Procedures

### List of Reagents

Chloramine-T trihydrate(Merck, Darmstadt, Germany), Copper(II)Sulphate, Penicillin, Streptomycin, RPMI-1640(Sigma-Aldrich, Burlington, MA, USA), RPMI-1640 (Euroclone, Pero, Italy), DMEM low glucose(Sigma-Aldrich), Trypsin-EDTA(Sigma-Aldrich), Blasticidin hydrochloride(Sigma-Aldrich), Trichloroacetic acid(Merck), Fetal Bovine Serum (Merck, Darmstadt, Germany), Iodine 125 Radionuclide (Perkin Elmer, Waltham, Mass., USA), Iodogen™ iodination reagent (Pierce, Thermo-Fischer), Alexa-488 succinimidyl ester (Thermo Fischer Scientific, Waltham, Mass., USA),, Anti-CD146 microbeads, anti-F4/80 microbeads (Miltenyi Biotech, Bergisch Gladbach, Germany), HSA Alburex (CSL Behring, King of Prussia, Penn., USA), Fetal Bovine Serum (Biowest, Nuialle, France), Resazurin (biotechne, Minneapolis, Minn., USA), Liberase™TM (Roche, Basel, Switzerland), Human fibronectin was extracted from human serum by affinity chromatography locally by the method of Vuento 1979^40^, Formaldehyde treated Serum Albumin (FSA) was prepared as described in Mego 1967^41^, Blomhoff 1984^32^, AGE-BSA was prepared as described in Hansen 2002^42^, Oxidized Low Density Lipoprotein (oxLDL) was prepared by Copper Sulphate oxidation as previously described in Li 2011^35^.

### Production and characterisation of oxHSA

Oxidation of HSA was carried out as described by Iwao 2006 ^24^; 300μM HSA was incubated with 100mM Chloramine-T in oxygen saturated PBS at 37°C for 1 hour. Afterwards the oxHSA was dialyzed against pure water and kept frozen until use. HPLC separation on Sperdex-200 10/300 (Amersham Pharmacia Biotech, Amersham, UK) size exclusion column was performed. Showing that the oxHSA eluted as three peaks of 837.4, 382 and 138.4 kDa (Figure S1).

### Radiolabeling of oxHSA, FSA, AGE-BSA, oxLDL

oxHSA or FSA was radiolabeled using carrier free 125-Iodine (Perkin-Elmer) according to the Iodogen™(Pierce) method and free iodine separated from protein by PD-10 (Cytiva) desalting column, as previously described Blomhoff 1984^32^. Specific activity was calculated from amount of added protein and measured activity post-labeling.

#### Animals

C57Black/6JRj mice were ordered from Janvier, and kept at the Department of Comparative Medicine, the Faculty of Health Sciences at UiT The Arctic University of Norway, under standard conditions with water and chow (SSniff, regular chow diet) ad libitum. Mice were between 8-14 weeks old for all of the procedures. All procedures were approved by the animal research authority under the food safety administration (Mattilsynet). Approvals FOTS ID: 30032, internal project number: 03/23 for in vivo component, internal project number: 09/22 for the in vitro component.

**In vivo clearance, organ- and hepatocellular distribution** was carried out as described in Santamaria-Simon 2014^43^. Briefly anesthetized mice were given intravenously 2-6μg ^125^I labelled oxHSA for biodistribution and hepatocellular distribution. For clearance blood samples were taken from the tail, in 2-5μL volumes over 30minutes, TCA precipitation was done to quantify intact/degraded ligand. For hepatocellular distribution animals were euthanized 5 min post-injection, and cells isolated as described in the section “Isolation of primary murine liver sinusoidal endothelial cells, Kupffer cells, hepatocytes”. Organ associated activities were measured on the Perkin-Elmer Wizard^2^, blood sample and isolated cell associated activities were measured on the Packard Cobra II auto-gamma.

### Isolation of primary mouse liver sinusoidal endothelial cells, Kupffer Cells, hepatocytes

Primary mouse LSEC, KC or HC were isolated as previously described in Elvevold 2022 ^44^. Briefly livers were perfused and digested with 1.2mg/50mL Liberase TM™(Roche) centrifuged to separate hepatocytes from non-parenchymal cell fraction, and followed by immune magnetic separation (MACS, Miltenyi) of LSEC and KC from the non-parenchymal fraction by CD-146 and F-4/80 respectively. Primary cells were cultured in serum-free RPMI-1640 (Euro-Clone/Sigma) supplemented with 10,000 U/mL Penicillin, 10 mg/mL Streptomycin, 1:100 (Sigma).

### HEK293 cells stably expressing stabilin 1 or 2

HEK293 cells expressing stabilin -1or -2, previously described in Hansen 2005^45^, or the empty vector pEF6V5His-TOPO (Merck) (plasmid containing Blasticidin resistance gene) were grown in DMEM low glucose (Sigma) supplemented with 10,000 U/mL Penicillin, 10 mg/mL Streptomycin, 1:100, (Sigma) 7% FBS (BioWest) and 10μg/mL Blasticidin hydrochloride (Merck)^45^.

### Affinity Chromatography

OxHSA, native HSA and FSA were coupled to cyanogen bromide activated Sepharose 4B (Pharmacia) as described in McCourt 1999^27^. Lysates from 19 million isolated LSEC were passed through the affinity columns, in 0.1% Triton TX-100 in PBS, columns were extensively washed with 0.1% TX-100/PBS and 0.1M Acetic acid pH 3, 0.01M EDTA.

Gel material was heated to 75^°^C in SDS, and sent for mass spectrometry analysis.

### Endocytosis experiments

Cells were seeded on human fibronectin-coated 48 well plates at 300K cells/ well for LSEC, 300 K cells for hepatocytes, 125-320K cells per well for KC depending on isolation yield, and allowed to adhere for 2 hours before use for LSEC, KC or 4 hours for hepatocytes, HEK cells were used after growing to confluence. For endocytosis experiments cells were kept in serum-free media with 1% native HSA (Alburex, CSL Behring) in RPMI-1640 (Euro-Clone) for LSEC, KC, hepatocytes and DMEM low glucose (Sigma) for HEK cells. Approximately 20000 cpm of labelled ligand, corresponding to approximately 5-15ng protein, was added to each well and cells were incubated for 2 hours (LSEC, KC, Hepatocytes) 4 hours (HEK cells) or a time course of 2, 4, 6, 18 hours (LSEC). After which cell associated, non-degraded and degraded fractions were collected and measured as described in Blomhoff 1984^32^. For competitive inhibition experiments several concentrations of non-radioactive ligand containing media were added to the cells immediately prior to addition of radiolabeled ligand. Iodine-125 measurements were done using the Cobra II auto-gamma (Packard).

### Fluorescent microscopy

oxHSA and FSA were labelled with Alexa488 using the manufacturer’s instructions (Thermo Fischer). Briefly labelling reagent was dissolved in DMSO and added to a 10mg/mL solution of protein in 0.1M bicarbonate buffer pH 8.3 for 1 hour at room temperature, then dialyzed against PBS in a 10K MWCO Slidealyzer dialysis cassette to remove uncoupled dye.

Cells were pre-stained with Cell Mask Orange (ThermoFischer)1:1000 for 5min, before addition of 10μg/mL Alexa-oxHSA for 30 minutes, after which cells were washed in PBS before being viewed under the EVOS (ThermoFischer) fluorescent light microscope.

### Scanning Electron Microscopy

Cells were seeded on human fibronectin covered glass 16 well plates at 25-40K cells/well and allowed to attach for 2 hours prior to treatment. Cells were treated with given concentrations of oxHSA in RPMI the indicated times and subsequently fixed with McDowell’s fixative. Cells were post-fixed with 1% OsO4 and dried with a graded series of ethanol (30, 60, 90, 100%) washes and finally hexamethyldisilane. Cells were sputter coated with Au/Pd immediately prior to scanning.

Scanning electron microscopy was performed the Zeiss Gemini or Sigma scanning electron microscopes at the advanced microscopy core facility at UiT.

### Viability experiments

Cell viability was assessed by LDH assay (Promega) or resazurin-resorufin (biotechne) assay were performed according to manufacturers instructions. For LDH LSEC were seeded 300K cells/well in a 48 well plate and treated with varying concentrations of oxHSA. After set timepoints the supernatants were collected and analyzed. For resazurin-resorufin cells were seeded the same way, with 1:10 resazurin reagent (biotechne) added to the culture media and measurements, using the ClarioStar plate reader wavelengths excitation 530-570nm emission 580-590nm, done at 3 and 6hours.

## Results

### In vivo biodistribution

The liver took up the majority of injected oxHSA of all the organs with on average 47% of total radiation, 6.5% GI-tract, 8% head, 2.6% kidneys, 2% tail, 1.3% spleen, 1.2% lungs, 1% heart and 30% remained in the carcass. The liver and spleen took up the most radioactivity per mass, 30% and 18% per gram respectively (Figure 1). The majority of the injected (75%) radiation (as estimated from injected dose or total radiation in organs+carcass) was cleared before the first blood sample was collected 0:55-1:40min post injection (Figure S2). Therefore, the t_1/2_ is even lower (<90sec).

**Figure 1:**
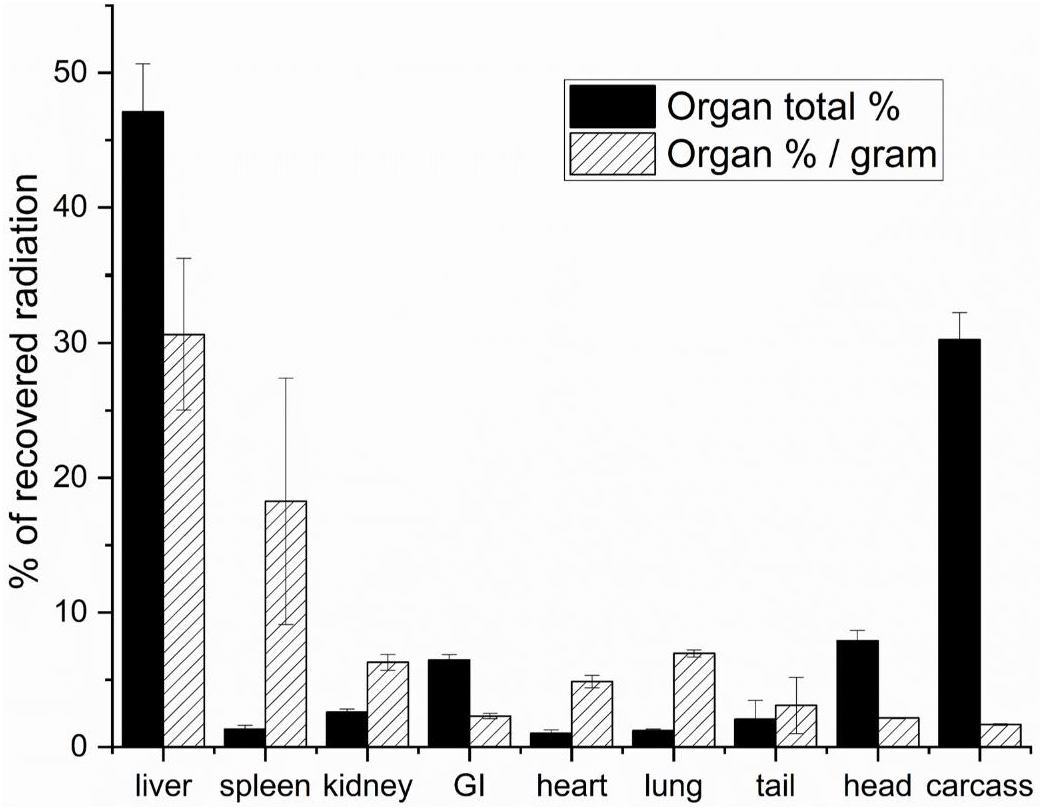
Biodistribution of oxHSA. 1-5μg 125I radio-labelled oxHSA was injected intravenously, and animals were sacrificed 30 min post-injection. Uptake is given as % of total recovered radioactivity (black bars) or as % of total recovered radioactivity per gram of organ (white/shaded bars). Results are given as averages ± standard deviation, n=3.

### Hepatocellular distribution

To determine the relative contribution of liver cells to oxHSA hepatocellular distribution was performed. Out of the cells of the liver the LSEC had the highest activities, compared to Kupffer cells or hepatocytes. LSEC contained activites (normalized to cell number) 15 and 11-fold higher than KC or hepatocytes respectively (Figure 2).

**Figure 2:**
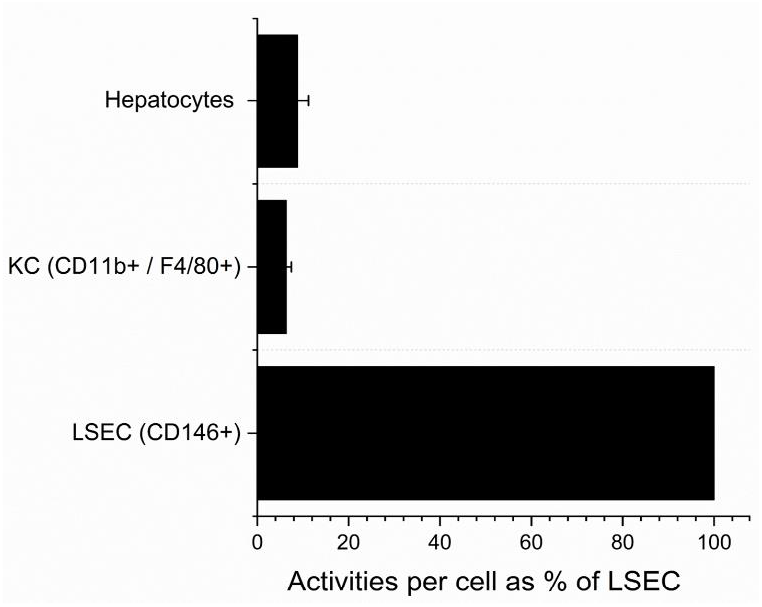
Hepatocellular distribution of oxHSA. Animals were injected with 1-5μg 125I radio-labelled oxHSA, sacrificed 5 minutes post-injection and LSEC, Kupffer cells and hepatocyte fractions were isolated. Graphs show radioactivity per cell normalized to LSEC, in isolated fractions of liver cells (selected by; CD146:LSEC, CD11b & F4/80:KC, Percoll 45%:hepatocytes). Results are given as averages ± standard deviation, n=3.

### In vitro identification of the oxHSA endocytosis receptor

LSEC detergent lysates were subjected to affinity chromatography on oxHSA coupled to Sepharose. A number of proteins were eluted from this column, including stabilins-1 and -2. Stabilins-1 and 2 were not eluted from control columns; i.e. Sepharose without protein, or Sepharose coupled with native HSA. Importantly, the cell lysates contained all cellular proteins, and not only cell-surface proteins.

Isolated murine LSEC showed the highest in vitro uptake with 35% uptake and degradation of added ^125^I-oxHSA over 2 hours of incubation increasing to 70% after 18 hours, per 300K cells (Figure 3A,B). Kupffer cells (resident macrophages) took up ≈ 13% of added ^125^I-oxHSA per 300K cells over 2 hours (Figure 3A). Hepatocytes took up ≈10% of added ^125^I-oxHSA per 300K cells over 2 hours, but this likely due to contamination by NPCs (Figure 3A).

**Figure 3:**
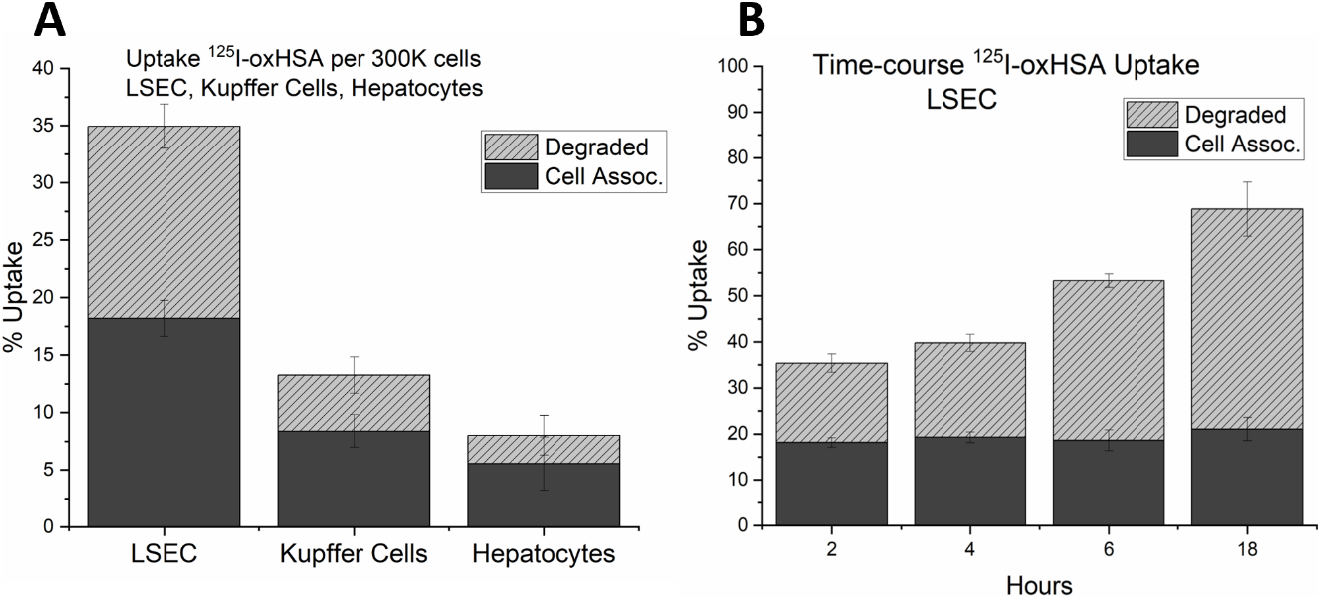
In vitro uptake of oxHSA by isolated liver cells. A) Uptake of 125I-oxHSA per 300K cells in LSEC, Kupffer cells and hepatocytes. LSEC and hepatocytes were seeded 300K/well, Kupffer cells were counted and uptake calculated per 300K cells. B) Time-course of 125I-oxHSA uptake in LSEC. Uptake is given as % of added (approx. 5-15ng/well). Solid bars indicate cell associated radioactivity, shaded bars indicate acid soluble radioactivity (= degraded ligand). Results are given as averages ± standard deviation, n=3.

To determine the potential role of the SR-H scavenger receptors stabilin-1 and stabilin-2, HEK293 cells stably over expressing mouse stabilin-1 and stabilin-2 were challenged with ^125^I-oxHSA. Both stabilin-1 and stabilin-2 HEK293 cells (but not the empty vector control) avidly endocytosed 48% and 67% of trace amounts of ^125^I-oxHSA, respectively, within 4 hours (Figure 4A-C). Figure 4A shows the % endocytosis of added ^125^I-oxHSA to the abovementioned HEK293 cells, as well as other known SR-H ligands: FSA; AGE-BSA and oxLDL. These other ligands were endocytosed at 29-32% and 32-55% by stabilin-1 and stabilin-2 HEK293 cells, respectively. The empty vector control cells endocytosed ≤12% of added ligand. The specificity of SR-H mediated uptake of oxHSA was tested by using oxHSA to inhibit uptake of other SR-H ligands. Stabilin-1 and stabilin-2 HEK293 cells were incubated with ^125^I-AGE-BSA (Figure 4B) or ^125^I-oxLDL (Figure 4 C) and challenged with unlabelled oxHSA (0-62 ug/ml or 0-5 ug/ml, respectively). ^125^I-AGE-BSA uptake was markedly inhibited in both SR-H expressing HEK293 cells at 7.5 ug/ml oxHSA. ^125^I-oxLDL uptake in the same cells was somewhat inhibited with 5.0 ug/ml oxHSA. Similar uptake and inhibition studies were performed on LSEC, which express both stabilin forms. LSEC challenged with 10μg/mL Alexa488-oxHSA for 30 min showed marked uptake as determined by fluorescent microscopy (Figure S3). AGE-BSA, FSA and oxLDL inhibited the LSEC uptake of ^125^I-oxHSA by 60-80% (Figure 4D). Unlabelled oxHSA inhibited the LSEC uptake of ^125^I-FSA (markedly), ^125^I-AGE-BSA (somewhat) and ^125^I-oxLDL (somewhat) (Figure 4E-G). Unlabelled oxHSA markedly inhibited LSEC uptake of ^125^I-oxHSA (Figure 4H), but not to the same degree as it did with FSA (Figure 4E).

**Figure 4:**
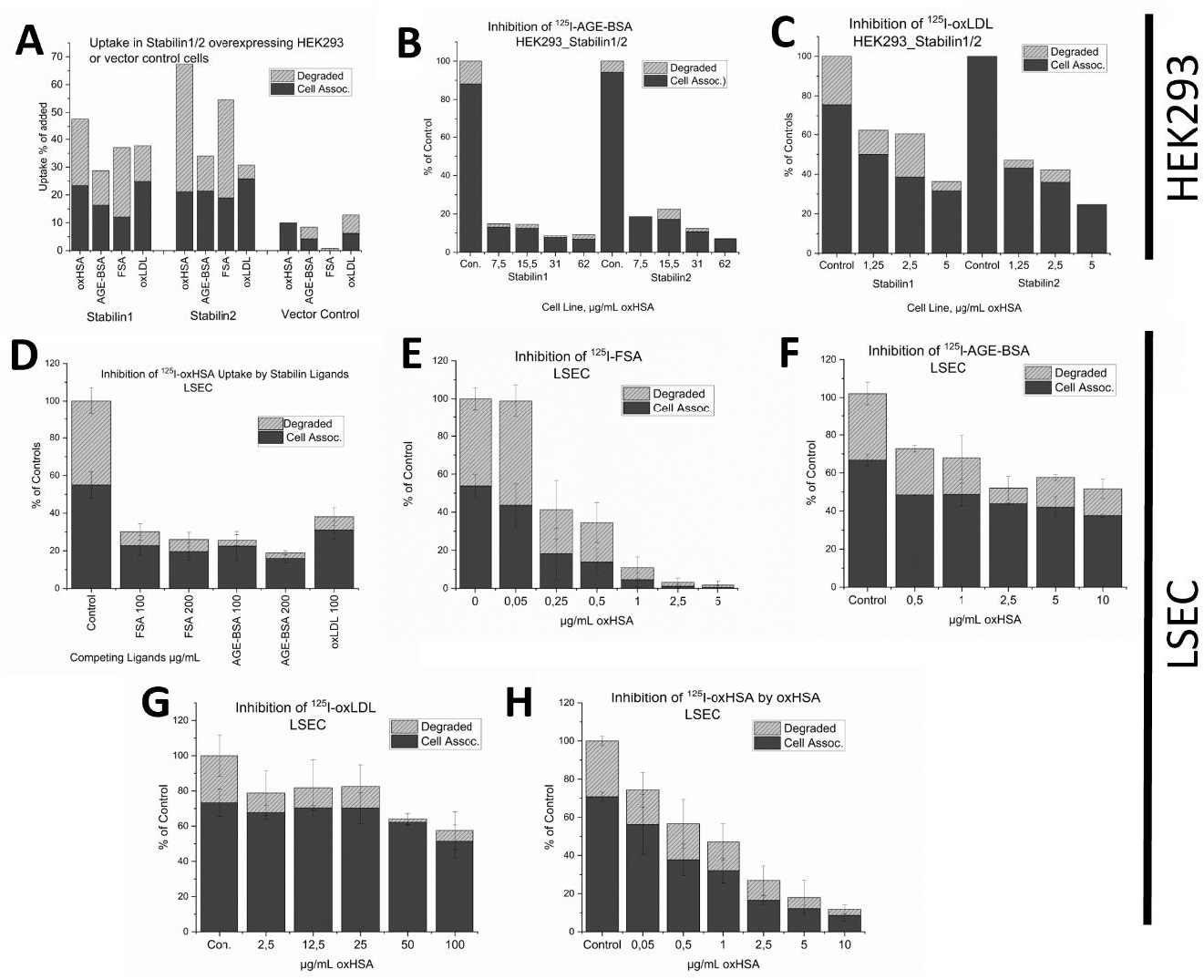
Stabilin -1 and -2 involvement in oxHSA uptake. A) Uptake of 125I-labelled oxHSA compared with uptake of other ligands for stabilin-1 and -2 (AGE-BSA, FSA, oxLDL) in HEK293 cells expressing stabilin -1, -2, or transfected with the empty vector. B) Inhibition of 125I-AGE-BSA uptake in stabilin-1 and -2 expressing HEK cells by oxHSA. C) Inhibition of 125I-oxLDL uptake in stabilin -1 and -2 expressing HEK cells by oxHSA. D) Inhibition of 125I-oxHSA uptake in LSEC by other ligands of stabilin -1 and -2 (FSA, AGE-BSA, oxLDL). E) Inhibition of 125I-FSA uptake in LSEC by oxHSA. F) Inhibition of 125I-AGE-BSA uptake in LSEC by oxHSA. G) Inhibition of 125I-oxLDL uptake in LSEC by oxHSA. H) Inhibition of 125I-oxHSA uptake in LSEC by unlabelled oxHSA. Uptake (in A) is given as % of radioactivity added per well, for inhibition graphs (B-H) uptake is given as % of (untreated) controls. Solid bars indicate cell associated radioactivity, shaded bars indicate acid soluble radioactivity (=degraded ligand). Results for LSEC are given as averages ± standard deviation, n=3.

To determine if the oxHSA-mediated inhibition of LSEC endocytosis was short or long term, we determined the level of FSA endocytosis after a 2-hour pulse of oxHSA (100 ug/ml) followed by chases of 3, 6 and 12 hours in RPMI media (Figure 5). There was little to no recovery of LSEC FSA endocytosis to control levels even after a 12-hour pulse of media (Figure 5) where levels were at 40% of untreated levels.

**Figure 5:**
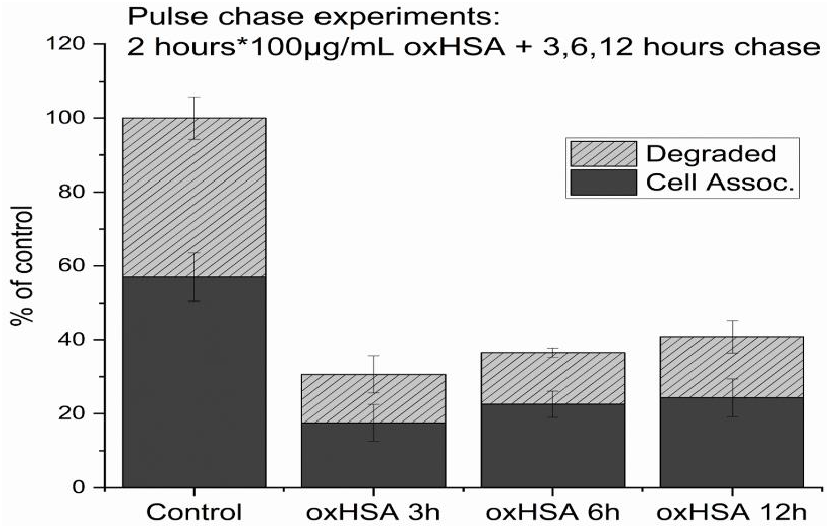
Pulse-chase / Recovery of endocytosis in LSEC. LSEC were treated with 100μg/mL oxHSA x 2hours, and then the indicated number (3, 6, or 12) of hours chase in cell culture media, before endocytosis experiments with 125I-FSA. Uptakes in % of matched untreated controls. Solid bars indicate cell associated radioactivity, shaded bars indicate acid soluble radioactivity (=degraded ligand). Results given as averages ± standard deviation, n=3.

### Morphology and viablity of LSEC challenged with oxHSA

LSEC treated with 10-160μg/mL oxHSA for 1 hours showed no morphological alterations at EM level (Figure S4). Cells treated with 0-320 μg/mL for 3-6 hours showed no changes to viability as measured by LDH or resazurin assays (Figure S5).

## Discussion

AOPP albumin, also known as oxHSA, is cleared from the circulation primarily by the liver and spleen ^24^. We synthesized oxHSA to determine the site of its uptake in the liver and its effect on liver cells. oxHSA characterization by HPLC revealed increased size peaks relative to HSA’s peak,(FigureS1) this is indicative of conformational rearrangement rather than added mass, as the electrophoretic motility under denaturing conditions (SDS) do not show such dramatic changes^24^. The HPLC profile of oxHSA is furthermore a very similar profile to model ligand FSA (data not shown). These conformational changes predispose albumin to scavenger receptor mediated clearance, judging by the examples of oxHSA and FSA. The oxHSA produced in this study was not toxic for LSEC as determined by LDH and resazurin assays, and the morphology of the cells was also seemingly unaffected by oxHSA as judged by SEM (Figure S4). We show that of all the liver cells, LSEC show the highest capacity for clearance of oxHSA (Figures 2 & 3). The most likely candidate receptors mediating this process are the SR-H scavenger receptors stabilin -1 and -2. This would be consistent with the observation that the highest stabilin expression levels are in the liver and spleen ^38^.

We established that oxHSA is cleared by stabilins -1 and -2 by uptake and competitive inhibition studies in LSEC and HEK293 cells constitutively expressing stabilin -1 and -2. For LSEC uptake of oxHSA was inhibited by FSA, AGE-BSA and oxLDL, and oxHSA in turn inhibited their uptake (Figure 4). The uptake of FSA was completely inhibited in LSEC by oxHSA, indicating a very similar binding profile. oxHSA moderately inhibited AGE-BSA in LSEC but inhibited AGE-BSA uptake very strongly in stabilin expressing HEK293 cells (Figure 4B,F). oxLDL uptake/degradation was slightly inhibited by oxHSA in LSEC but very strongly in stabilin expressing HEK293 cells (Figure 4C,G), suggesting these ligands (AGE-BSA, oxLDL) have additional receptors for endocytosis in LSEC.

We performed pulse chase experiments to determine if the effect of oxHSA on endocytosis was long lasting. Endocytosis was reduced to 40% of controls 12 hours after challenge with 100μg/mL for 2 hours (Figure 5), which is comparable to previously described effects of AGE-BSA on endocytosis mediated by stabilins -1 and -2^42^. This suggests that oxHSA depletes binding activity over a physiologically relevant timeframe. Thus, circulating oxHSA may impair the clearance of other stabilin ligands, which may be of concern during pathological states with high oxidative stress. For example it has been shown that stabilins in the liver are responsible for the elimination of LPS arriving from the gut, preventing systemic inflammation ^46^. It has previously been shown that SR-H deficiency causes kidney fibrosis in a mouse model ^37^. This was also suggested as a link between diabetic AGE formation and diabetic reno-pathy, which also sees heightened levels of oxidation protein products ^35^. Partial hepatectomy often leads to kidney injuries ^47^, where a reduction in clearance of scavenger receptor ligands may be a driver of these injuries. This fits with the presence of oxidized albumin in uremic patients ^24^ as either a marker for reduced clearance or a uremic toxicant itself.

Additionally, it has been shown that the scavenger endothelium of the liver is the main site of clearance for pro-atherogenic molecules such as oxidized LDL, and AGEs ^35,48^. An increased circulation time, or accumulation of these ligands is likely to cause atherosclerotic plaques and localized inflammation in the vasculature. Oxidative stress and detection of oxidation protein products has been linked with atherosclerosis previously ^4^, with AOPP-Albumin been shown to cause atherosclerotic plaque formation in rabbits ^49^.

This suggests a common theme and possible feedback mechanism for these ligands, where an increase above a threshold will lead to a vicious cycle, where AOPP clearance is inhibited by their own prevalence, and their prevalence induces their own formation by an oxidative stress/ inflammation related mechanism at the sites of deposition. Thus, atherosclerosis and systemic inflammation is both driving and being driven by AOPP formation. This would all have implications for other organs, such as kidneys, as suggested by Schledzewski et al. ^37^.

Similarly, the pathogenic progression of liver disease or injury, would lead to a reduction in clearance of the atherogenic oxidation protein products (oxHSA, oxLDL etc.) as was indeed found by Öettl in 2013^14^, which would increase their relative concentrations, circulation time, leading to deposition, plaque formation and inflammation. Plausible mechanisms driving this would be the impaired synthesis of new albumin by hepatocytes coupled with impaired clearance of modified albumins from circulation by LSECs.

In summary oxHSA is cleared in vivo by the LSEC, is a ligand for stabilins -1 and -2, and *in vitro* challenge of LSEC with oxHSA causes downregulation of SR-H mediated endocytosis. This has implications for the clearance of waste proteins, LPS and other ligands normally cleared by SR-H, since elevated levels of oxidized albumin are seen in diseases such as atherosclerosis, diabetes and acute and chronic liver failure. If oxidized albumin interferes with SR-H mediated clearance, this may explain some of the downstream effects of pathological inflammation. Strategies inhibiting the formation of oxidation protein products during disease and inflammation may thus be warranted. Interventions such as those reviewed by Forman & Zhang 2021^50^ may be of use in such cases.

## List of Abbreviations

AGE: advanced glycation end products
AOPP: advanced oxidation protein products
BSA: bovine serum albumin
FSA: formaldehyde modified BSA
HDL: high density lipoprotein
HPLC: high performance liquid chromatography
HSA: human serum albumin
KC: Kupffer cell(s)
LDH: lactate dehydrogenase
LDL: low density lipoprotein
LPS: lipopolysaccharide
LSEC: liver sinusoidal endothelial cell(s)
MWCO: molecular weight cut-off
oxLDL: oxidized LDL
oxHSA: oxidized HSA
PBS: phosphate buffered saline
SDS: sodium dodecyl sulphate
SR-H: scavenger receptor class
H, t_1/2_: half-life

## Acknowledgements

Special thanks to Randi Olsen and Tom Ivar Eilertsen at the microscopy core facility, Jack Ansgar Bruun at the UiT proteomics facility, Montserrat Martin-Armas at the PET-CORE centre at UiT, Bård Smedsrød, Karen K. Sørensen, Cristina I. Øie, for advice on practical details on endocytosis assays and application forms.

## Supplemental Figures

**Figure S1:**
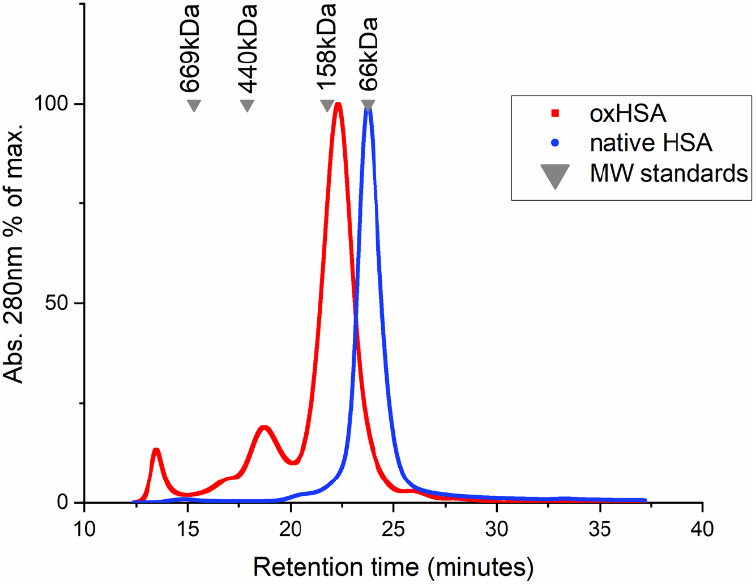
**Supedex-200 10/300 HPLC chromatogram** of oxHSA (red) superimposed on chromatogram of native HSA (blue), MW standards are indicated with gray triangles. X-axis: retention in minutes. Y-axis: absorbance at 280nm as % of maximum.

**Figure S2:**
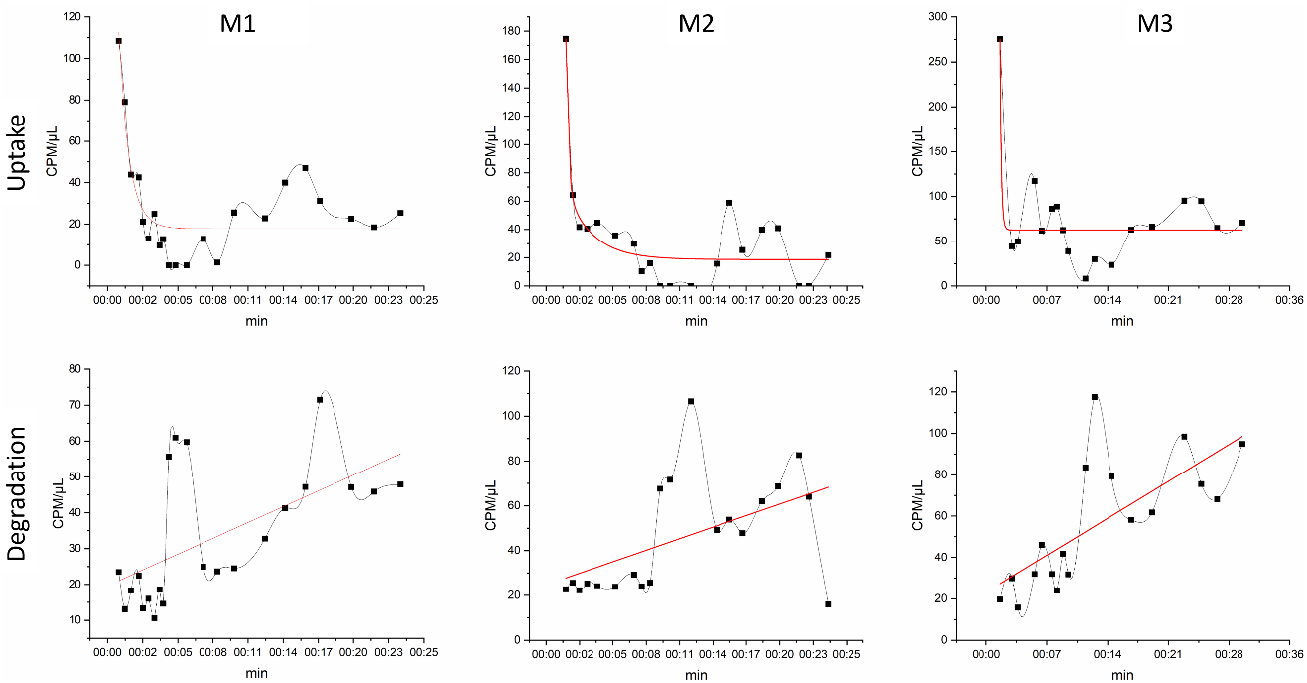
**Blood clearance curves** from biodistribution study. X-axis: time in minutes, y-axis: CPM/μl blood. 3 mice were injected with 1-5μg ^125^I labelled oxHSA, and blood samples collected 0-30minutes post-injection, each animal is plotted separately. Top panels: Removal of acid insoluble (intact protein) radiation from blood. Bottom panels: appearance of acid soluble radiation (degraded protein).

**Figure S3:**
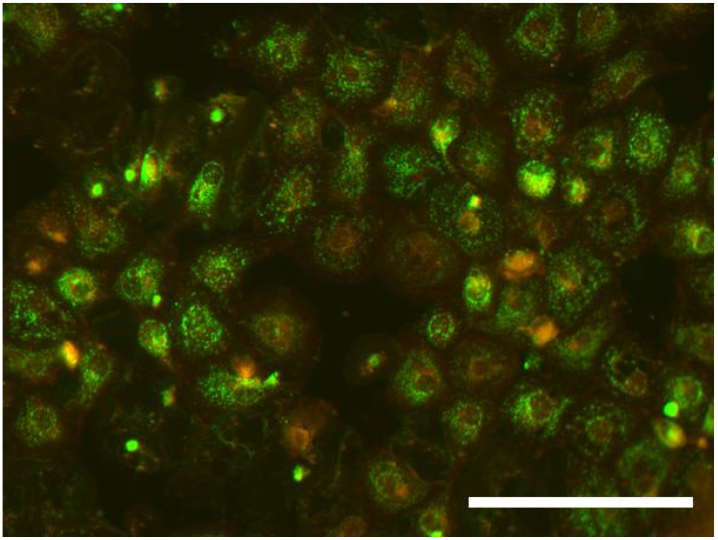
**Fluorescence micrograph** of LSEC given Alexa488 labelled oxHSA (10μg/mL x 30min). Cell membranes were stained with Cell Mask Orange 1:1000 × 5 min before addition of oxHSA. oxHSA staining shows expected vesicular pattern. Scale bar = 100μm.

**Figure S4:**
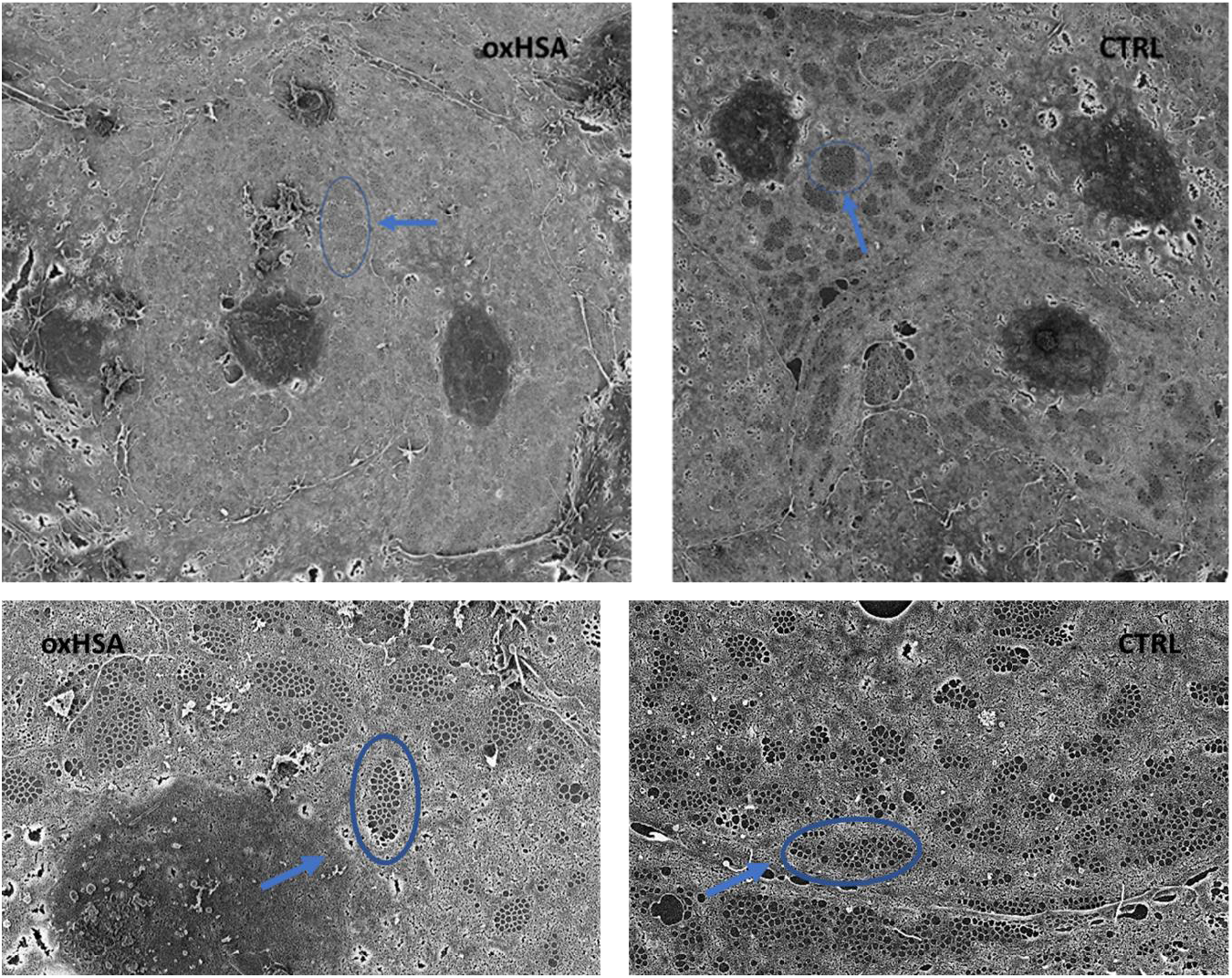
**Scanning Electron Micrographs** of LSEC treated with 160μg/mL oxHSA (**oxHSA**) and untreated controls (**CTRL**). Representative images, overview/large FOV (top) or close-up (bottom), of LSEC treated with 0 (CTRL) or 160μg/mL oxHSA (oxHSA) for 1 hour. Arrows and ellipses indicate fenestrations organized into sieve-plates.

**Figure S5:**
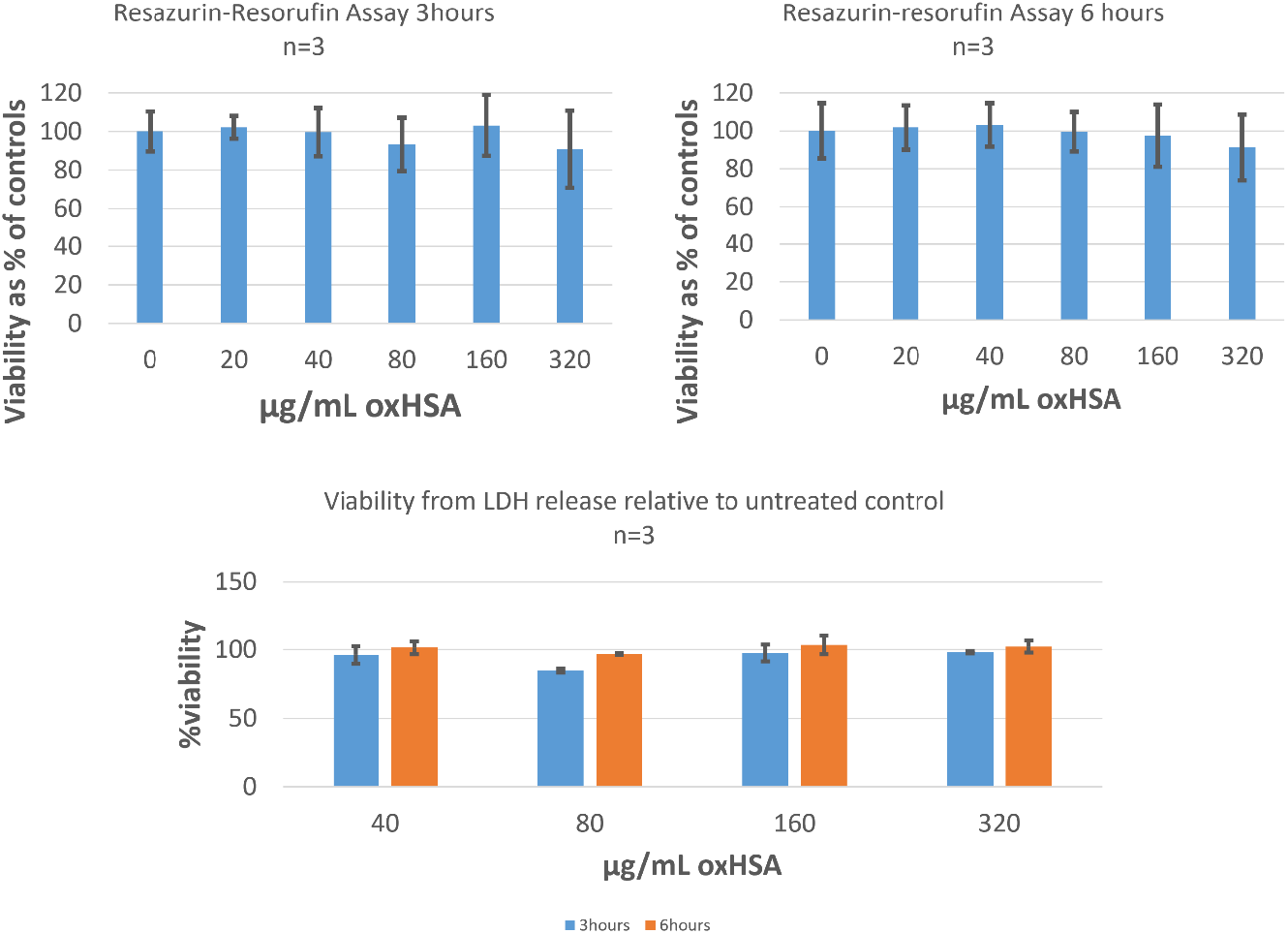
**Viability assays** performed on LSEC. TOP PANNELS Resazurin-resorufin assay: 300K cells were treated with indicated concentrations of oxHSA with resazurin added to the culture media, measurements were done at 3 and 6 hours after addition of oxHSA. BOTTOM PANELS LDH-Glo assay: 300K cells were treated with indicated concentrations of oxHSA, and supernatant samples collected at 3 and 6 hours. Viability calculated from LDH release relative to positive control (triton x-100) and negative control (untreated cells).

